# OprF functions as a latch to direct Outer Membrane Vesicle release in *Pseudomonas aeruginosa*

**DOI:** 10.1101/2023.11.12.566662

**Authors:** Shrestha Mathur, Susan K. Erickson, Leah R. Goldberg, Sonia Hills, Abigail G.B. Radin, Jeffrey W. Schertzer

## Abstract

Bacterial Outer Membrane Vesicles (OMVs) contribute to virulence, competition, immune avoidance and communication. This has led to great interest in how they are formed. To date, investigation has focused almost exclusively on what controls the initiation of OMV biogenesis. Regardless of the mechanism of initiation, all species face a similar challenge before an OMV can be released: How does the OM detach from the underlying peptidoglycan (PG) in regions that will ultimately bulge and then vesiculate? The OmpA family of OM proteins (OprF in *P. aeruginosa*) is widely conserved and unusually abundant in OMVs across species considering their major role in PG attachment. OmpA homologs also have the interesting ability to adopt both PG-bound (two-domain) and PG-released (one-domain) conformations. Using targeted deletion of the PG-binding domain we showed that loss of cell wall association, and not general membrane destabilization, is responsible for hypervesiculation in OprF-modified strains. We therefore propose that OprF functions as a ‘latch’, capable of releasing PG in regions destined to become OMVs. To test this hypothesis, we developed a protocol to assess OprF conformation in live cells and purified OMVs. While >90% of OprF proteins exist in the two-domain conformation in the OM of cells, we show that the majority of OprF in OMVs is present in the one-domain conformation. With this work, we take some of the first steps in characterizing late-stage OMV biogenesis and identify a family of proteins whose critical role can be explained by their unique ability to fold into two distinct conformations.

**Significance:** Vesicular transport is now recognized to operate in all domains of life. However, the study of OMV biogenesis has been challenging because genetic screens failed to identify proteins analogous to those involved in eukaryotic vesicular transport. With this work we identify the first protein whose direct action can both define the location and govern the mechanism of OMV release. Our latch model is consistent with previous observations linking OmpA family proteins to OMV biogenesis, but further describes a physical mechanism that has broad implications for vesicle production and function across species. The work presented here advances our understanding of a fundamental virulence-associated process in bacteria, while underscoring stark differences in how transport vesicles are formed in prokaryotes *versus* eukaryotes.

## Introduction

The production of extracellular vesicles derived from the Outer Membrane (OM) of Gram-negative bacteria is now recognized to be a ubiquitous phenomenon. These Outer Membrane Vesicles (OMVs) range in size from 50-300nm in diameter, are comprised of OM components released from the parent cell, and have been discovered to transport diverse and seemingly regulated cargo (1–8). In fact, vesicular transport has been recognized as a distinct secretion system within the bacterial toolbox (1, 9). The ability of OMVs to transport cargo at high concentration over long distances while protected from the environment helps to explain their prominent role in many important bacterial processes. These include virulence (1, 2, 10–13), interbacterial competition (14–16), immune modulation and avoidance (3, 17–21), horizontal gene transfer (12, 22–24), cell-cell communication (25–27), nutrient acquisition (7, 28) and biofilm development (29–32).

Work in recent years has provided insight into how OMVs are formed, with several models proposed to describe the process in different species. Detailed mechanisms have been described that involve peptidoglycan (PG) homeostasis (33–36), stress response (37–44), manipulation of membrane lipid composition (8, 45–49), intercalation of membrane-active small molecules (27, 42, 50–57), and outright cellular explosion (58, 59). Understanding gained from previous work has focused largely on the initiation of OMV biogenesis, particularly on how the membrane is induced to bulge to begin the process. With this work, we set out to explore later stages in OMV development, specifically the requirement that regions of the OM intended for release as OMVs must become detached from the underlying PG layer. OMV production has loosely been associated with the idea of loss of OM-PG linkage for some time (35, 60–68). However, the nature of investigation in this area has often made it difficult to distinguish between effects on natural OMV biogenesis pathways vs. unrelated membrane shedding or cellular disintegration.

Proteins that anchor the OM to the PG tend to be excluded from the cargo of OMVs (69, 70). Interestingly, this is not true for one family of highly conserved OM-PG linkage proteins. Although the OmpA family of proteins has a well-established role in OM-PG linkage (60, 71), members of this group are found to be abundant in the OMVs of several species (72–78). We reasoned that this discrepancy might be explained by the unique ability of these proteins to adopt two distinct conformations. OmpA, and its homolog OprF in *P. aeruginosa*, are examples of “slow porins” found in the OM of a wide variety of Gram-negative species. These proteins are capable of folding into a beta-barrel structure that produces a large diameter pore (like a classic porin) but displays remarkably low permeability when reconstituted into liposomes or planar bilayers (reviewed in (79)). These seemingly contradictory observations are reconciled by the fact that OmpA homologs can naturally fold into two conformers: a “closed” two-domain conformation where the C-terminal region folds into a globular domain in the periplasm and interacts with PG, and an “open” one-domain conformation where the C-terminal region joins the N-terminal region embedded in the membrane to produce the large beta-barrel pore. Studies in *E. coli* and *P. aeruginosa* have determined that >90% of OmpA and OprF found in the OM exist in the “closed” two-domain conformation, explaining the major role of these proteins in OM-PG linkage as well as the low permeability of solutes through these proteins in cells and reconstituted systems (79–82).

In this study, we test the hypothesis that OprF in *P. aeruginosa* helps control OMV production through its ability to exist in two different conformations. First, we demonstrate that the specific loss of PG-interaction ability explains hypervesiculation in *oprF* mutant strains, rather than general membrane instability caused by the loss of this abundant OM protein. We then develop a methodology to measure the conformational distribution of OprF in cellular membranes vs. OMVs and use this technique to discover that the ratio of the one- to two-domain conformers is inverted in OMVs relative to cells, with the vast majority in OMVs adopting the one-domain structure. We propose that OprF functions as a “latch” to direct OMV release by reducing the local density of OM-PG linkages in regions of the OM destined to become OMVs.

## Results

### OprF variants properly translocate to the OM and correctly fold

To test our hypothesis that OM-PG connections mediated by OprF are important for regulating OMV formation, we first created constructs to express two variant proteins: OprF ΔN_265_-V_276_ (OprF-PG_del_) is missing the majority of its PG-association motif (83, 84), while OprF ΔK_164_-K_326_ (OprF-C_del_) is missing the entire C-terminal domain (which contains the PG-association motif). Before testing whether either variant could rescue the Δ*oprF* hypervesiculation phenotype (67), we needed to confirm that the variant proteins were properly translocated to the OM and properly folded. Each variant (including native OprF) was expressed in *oprF* null mutant PA14 and PAO1 strains and subcellular localization was analyzed by Western blot using MA7-1 monoclonal antibody specific to the N-terminal region of OprF (85) (kind gift of Dr. R.E.W. Hancock) following cellular disruption and subcellular membrane fractionation by sucrose density centrifugation. Before analysis, successful membrane separation was confirmed by probing each fraction for KDO sugar (OM marker) and succinate dehydrogenase activity (IM marker) (see Fig S1). Fig 1 A-B shows that the vast majority of each OprF variant is found in the OM of both the PA14 and PAO1 mutant strains. Representative raw Western blots are presented in Fig S2. Expression of the OprF variants was also confirmed to have no effect on cell growth (Fig S3A-B).

**Fig 1.**
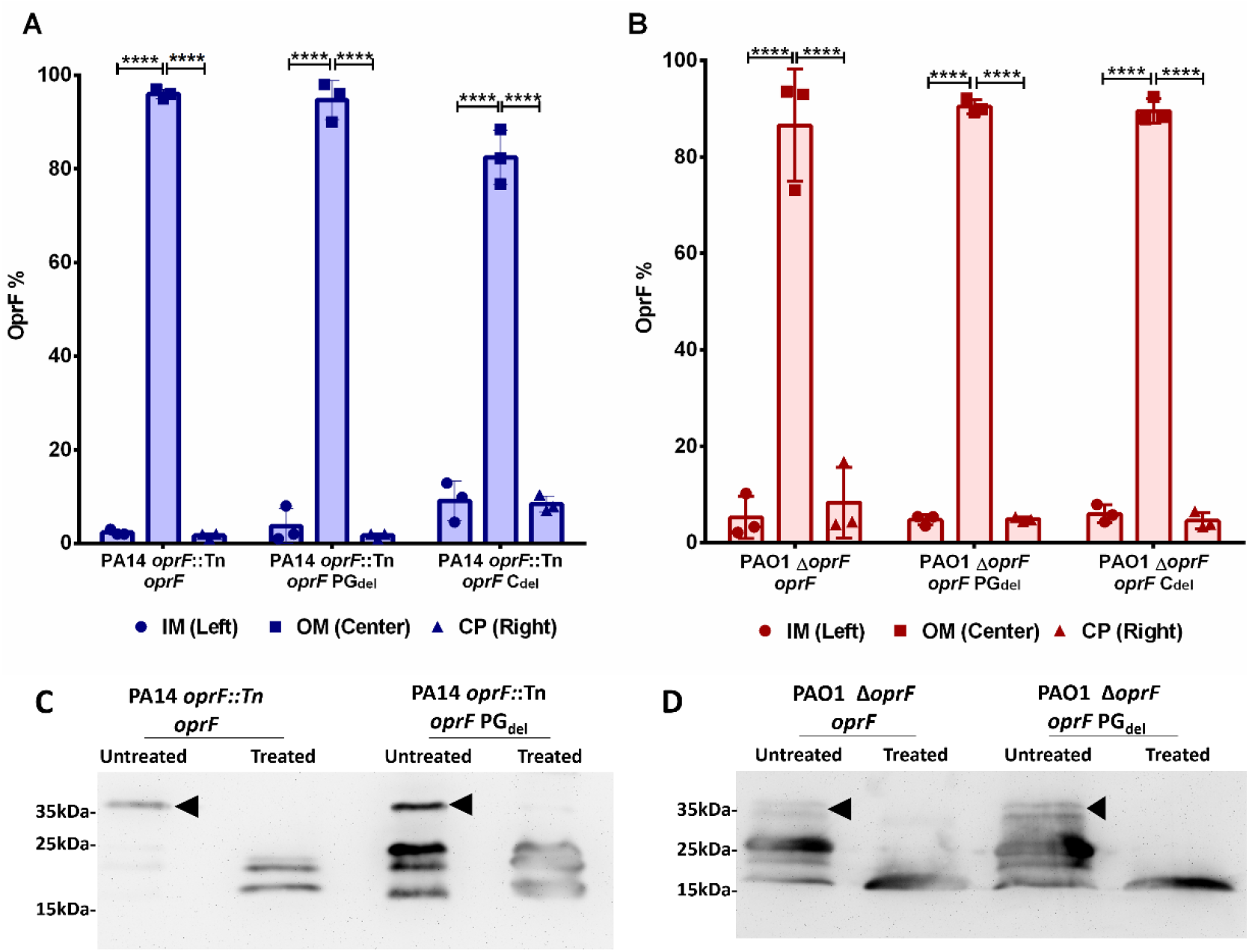
Validation of OprF variant translocation and conformation in the OM. The proportion of OprF variant proteins present in the inner membrane (IM), outer membrane (OM), and cytoplasm (CP) after expression in the PA14 (A) and PAO1 (B) *oprF* null strains was assessed by Western blot (also see Fig S2). Error bars represent standard deviation. Statistical significance was analyzed by Two-way ANOVA followed by Tukey’s multiple comparisons test (****p < 0.0001; n = 3). Purified OM fractions from the PA14 (C) and PAO1 (D) *oprF* null strains expressing full length or PG_del_ OprF variants were subjected to proteolysis by Proteinase K and analyzed by Western blot using N-terminal specific monoclonal antibody. The full length OprF is marked by black arrows in the untreated lanes, while degraded OprF fragments are seen between 15-20 kDa after treatment with proteinase K. Blots are representative of three independent experiments.

To assess proper folding of the OprF variant proteins, we employed a proteolysis protection assay performed on purified OM from *oprF* null mutant strains expressing each OprF variant. It is reported that 95% of OprF in the OM adopts the two-domain conformation in wild type cells (82). Proteinase K treatment of purified OM fragments would therefore be expected to degrade the exposed C-terminal domain while leaving the OM-embedded N-terminal domain intact. Degradation of the C-terminal domain after proteinase K treatment was analyzed by Western blot using MA7-1 antibody. Since OprF-C_del_ is missing the entire C-terminal domain and previous reports have confirmed that it folds correctly when expressed *in trans* (86), this analysis was only performed for native OprF and OprF-PG_del_. Fig 1 C-D confirms that native OprF overwhelmingly adopts the two-domain conformation in the OM and that the C-terminal domain is selectively degraded while the N-terminal domain is left intact. Importantly, the pattern is the same for the OprF-PG_del_ variant.

### OprF variants fail to rescue the Δ*oprF* hypervesiculation phenotype

Removal of the entire OprF protein results in a hypervesiculation phenotype (67). However, complete loss of such an abundant and important protein may result in unintended membrane stress phenotypes that could confound anaylsis of OMV formation. To address this, we used targeted OprF sequence variants to address whether the specific loss of OprF’s ability to interact with PG was responsible for hypervesiculation. Fig 2 A-B shows that expression of native OprF in both the PA14 and PAO1 null mutant backgrounds resulted in complete rescue of the hypervesiculation phenotype. Notably, both the OprF-PG_del_ and OprF-C_del_ variants failed to rescue the hypervesiculation phenotype, with OprF-C_del_ being indistinguishable from the *oprF* null mutant and OprF-PG_del_ showing partially reduced hypervesiculation.

**Fig 2.**
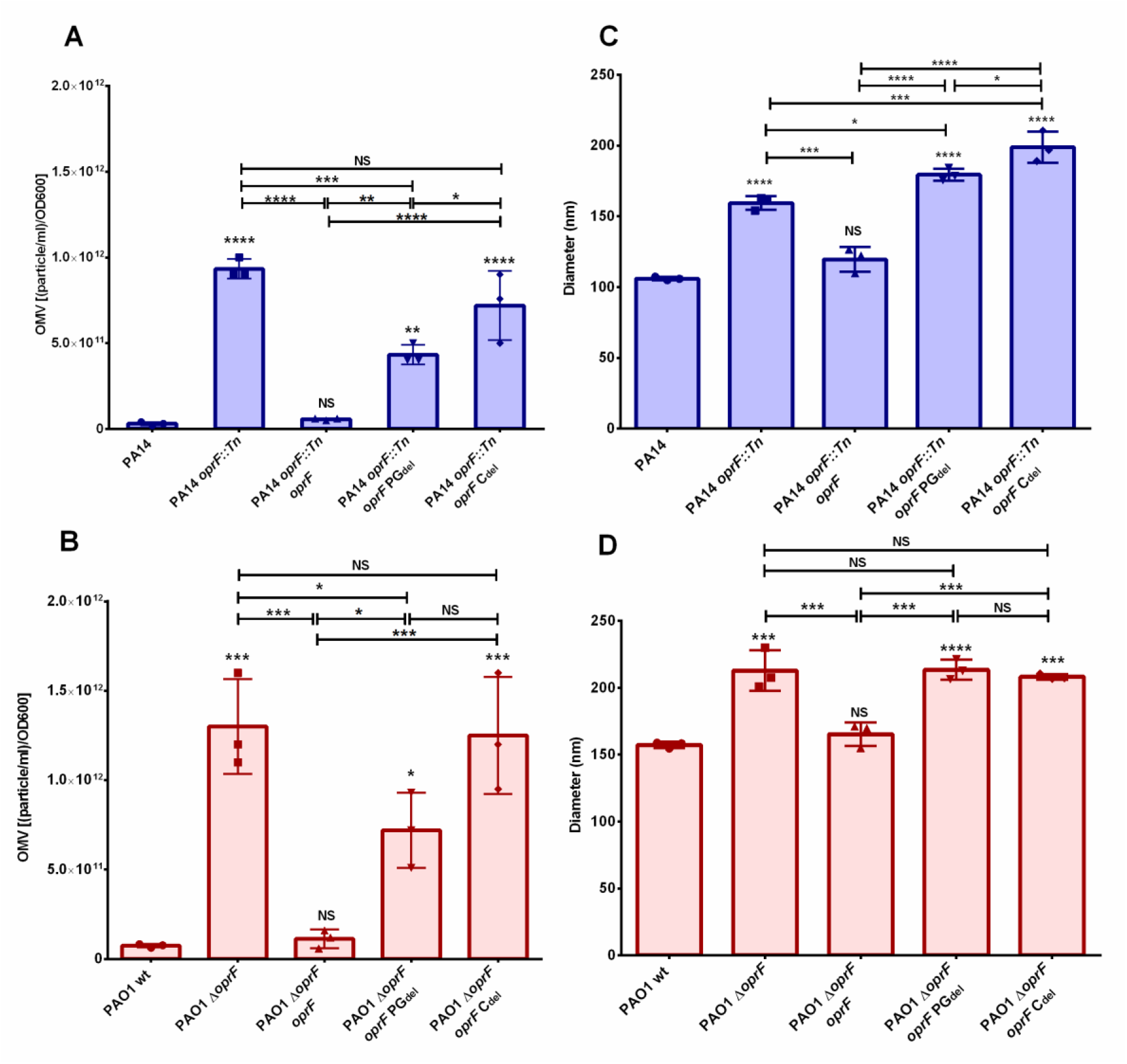
Analysis of OMV quantity and size. OMVs were harvested from the identified strains and analyzed by Nanoparticle Tracking Analysis (NTA). Vesicle quantity is presented in panels A & B. The mode of the vesicle size distribution is presented in panels C & D. Error bars represent standard deviation. Statistical significance was analyzed by One-way ANOVA followed by Tukey’s multiple comparisons test (* p < 0.05; ** p < 0.01; *** p < 0.001; ****p < 0.0001, NS = not significant; n = 3). Asterisks directly over bars denote differences from WT; asterisks associated with horizontal lines refer to differences between the indicated bars.

Analysis of OMV production using Nanoparticle Tracking Analysis (NTA) provided us information about the size distribution and quantity of vesicles in each sample. This allowed us to discover that the mode diameter of the OMVs produced by the *oprF* null mutants was significantly larger than wild type. The pattern of rescue of this phenotype by expression of native or variant OprF proteins was identical to that for the hypervesiculation phenotype, suggesting that the ability of OprF to attach to PG is involved in controlling OMV size as well as the level of OMV production (Fig 2 C-D).

### Hypervesiculation in OprF variant strains is not due to general membrane instability or cell lysis

To ensure that the hypervesiculation phenotypes observed were not the result of membrane instability arising from alteration of the OprF protein, all strains were challenged to grow on LB agar supplemented with 0.1% sodium deoxycholate. CFUs recovered in the presence of detergent were compared to CFUs recovered in the absence of detergent as a measure of general membrane stability. Fig 3 A-B shows that the reduction in viability upon exposure to deoxycholate was not significantly different from wild type for any of the mutant strains, eliminating the possibility that gross membrane instability was responsible for the hypervesiculation phenotypes observed. Similarly, we analyzed the purified OMVs for the presence of succinate dehydrogenase enzyme activity. Succinate dehydrogenase is an integral inner membrane protein, and its presence in OMV preps would provide evidence of cell lysis. Fig 3 C-D shows that no succinate dehydrogenase activity was detected in the OMV preps from any of the strains tested, demonstrating that hypervesiculation was not due to any vesicle production mechanism that involves cellular disintegration or explosive lysis (58, 59).

**Fig 3.**
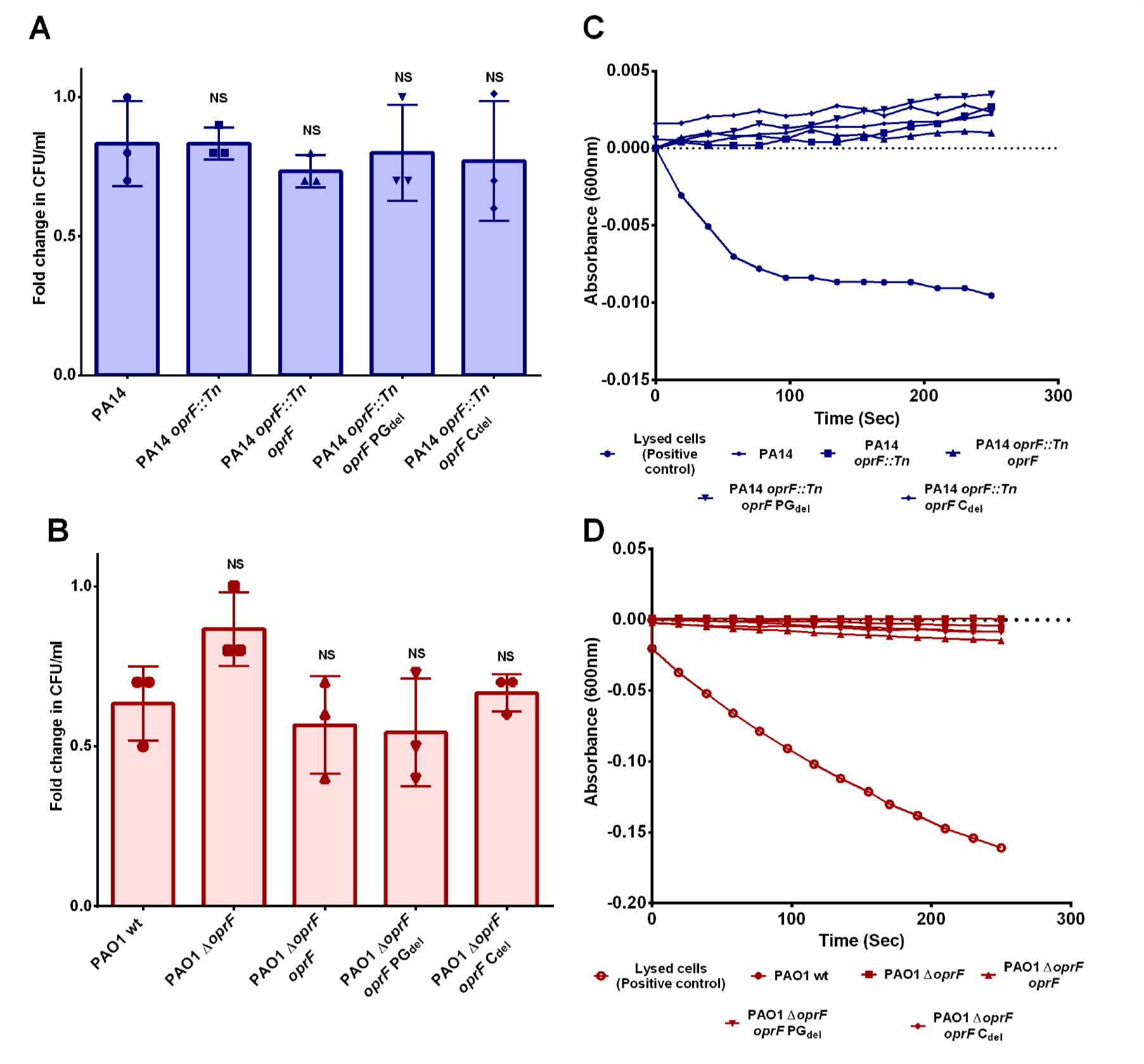
Analysis of membrane stability. Susceptibility to 0.1 % sodium deoxycholate was tested for all the strains in both the PA14 (A) and PAO1 (B) backgrounds. Error bars represent standard deviation. Statistical significance was analyzed by One-way ANOVA followed by Dunnett’s multiple comparisons test (NS = not significant; n = 3). Succinate dehydrogenase (SDH) enzyme activity was measured in OMVs prepared from all strains in both the PA14 (C) and PAO1 (D) backgrounds and compared to lysed bacterial cells (positive control). SDH assay results are representative of three independent experiments.

### Hypervesiculation in OprF variant strains is not due to overproduction of PQS

Loss of the entire OprF protein was previously shown to result in hypervesiculation, but the direct attribution of mechanism was confounded by an accompanying increase in PQS production in the *oprF* mutant strain (67). We directly measured PQS production in each of our strains using thin layer chromatography. Fig 4 shows that there was no increase in PQS production for any strain in our system. PQS production in PA14 strains was indistinguishable from one another. In PAO1, all mutant strains unexpectedly produced less PQS than wild type, though still within the typical physiological range. A *decrease* in PQS production can not explain hypervesiculation as a result of small molecule induced OMV biogenesis in these strains. Rather, these results are consistent with hypervesiculation being the result of the loss of specific interactions between OprF and PG.

**Fig 4.**
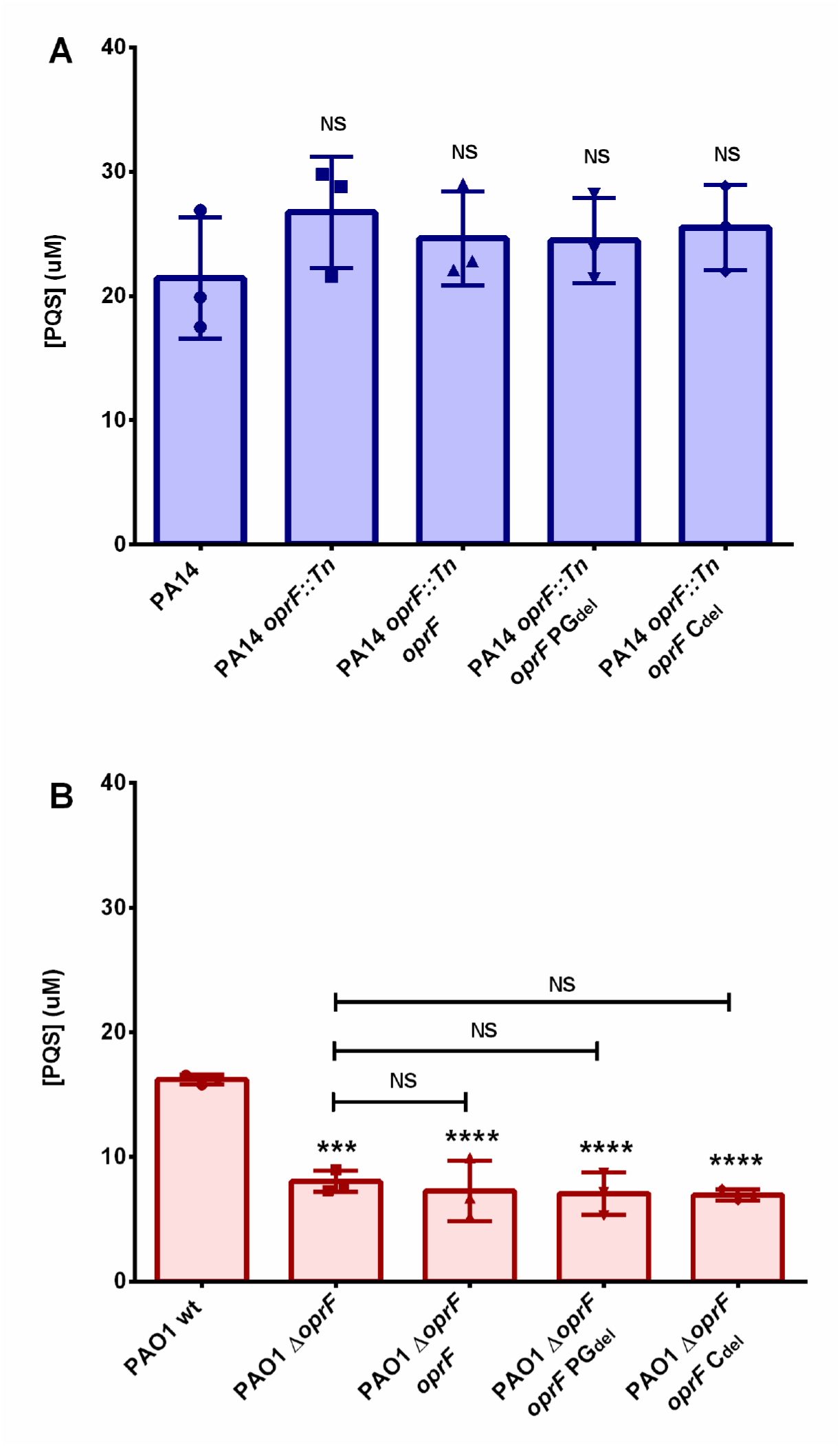
Analysis of PQS production. PQS was extracted from all strains in both the PA14 (A) and PAO1 (B) backgrounds and quantified using thin layer chromatography. Error bars represent standard deviation. Statistical significance was analyzed by One-way ANOVA followed by Dunnett’s or Tukey’s multiple comparisons test (*** p < 0.001; ****p < 0.0001, NS = not significant; n = 3). Asterisks directly over bars denote differences from WT; notations associated with horizontal lines refer to differences between the indicated bars.

### OprF is equally sorted between the OM and OMVs

To investigate the conformational distribution of OprF in OMVs *vs.* cells, we constructed an epitope tagged variant. A small 9 amino acid V5 epitope tag was inserted at position 312 in the C-terminal region of the protein. This position was chosen to correspond with previous strategies used to topologically map OprF (82, 85). From those studies, position 312 is known to be protected in the periplasm in the two-domain conformer and exposed on the cell surface in the one-domain conformer. To ensure antibody access to the V5 epitope on the surface of cells and OMVs, we expressed the OprF-V5 variant in the O-antigen-lacking Δ*wbpL* background. The presence or absence of O-antigen has been shown to have no effect on the level of OMV production (87). As was done for our previous OprF variants, we first established that OprF-V5 was properly translocated to the OM (Fig 5A, Fig S4) and its expression did not affect growth, membrane stability, OMV number & size, or PQS production (Fig S5).

**Fig 5.**
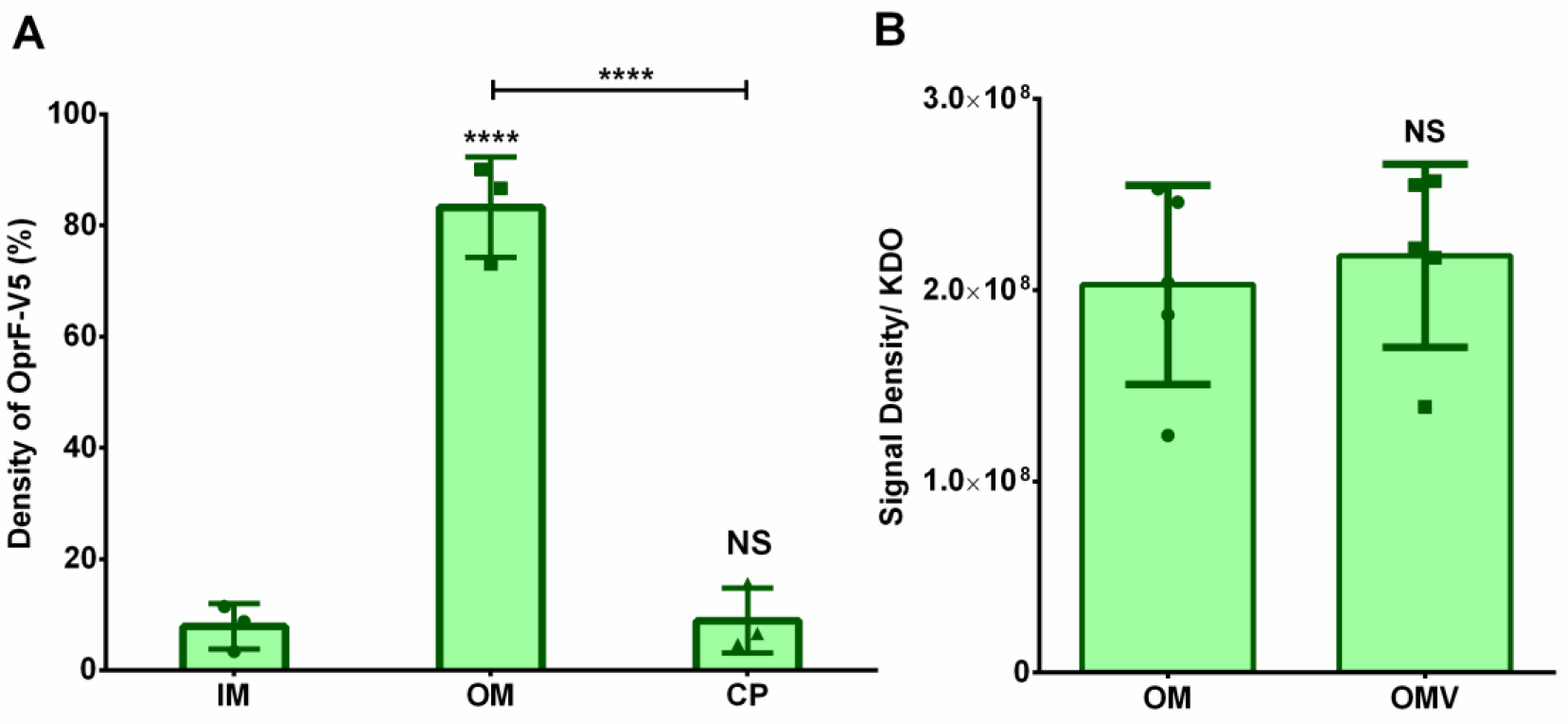
Translocation and sorting of OprF-V5 between the OM and OMVs. (A) Proportion of OprF-V5 found in the inner membrane (IM), outer membrane (OM), and cytoplasm (CP) when expressed in PAO1 Δ*wbpL oprF* V5 (also see Fig S4). Error bars represent standard deviation. Statistical significance was analyzed by One-way ANNOVA followed by Tukey’s multiple comparisons test (****p < 0.0001, NS = not significant; n = 3). (B) The amount of OprF-V5 present per unit membrane in OM *vs*. OMVs was assessed by normalizing anti-V5 Western blot signal to the concentration of KDO in each fraction (also see Fig S6). Error bars represent standard deviation. Statistical significance was analyzed by Two-tailed unpaired t-test with Welch’s correction (NS = not significant; n = 5). Asterisks directly over bars denote differences from WT; asterisks associated with horizontal lines refer to differences between the indicated bars.

Following this, it was important to understand whether OprF is preferentially sorted into or excluded from OMVs. To accomplish this, OMVs were purified and the amount of OprF-V5 was quantified by Western blot and normalized to the amount of KDO sugar in the sample. The same procedure was performed on purified OM from disrupted and density fractionated cells. Comparing the KDO-normalized OprF-V5 signal in each of these samples allowed us to assess how much OprF was present “per unit membrane”. Fig 5B shows that OprF density is equivalent in these two fractions, demonstrating that there is no preferential sorting of OprF between OMVs and the OM. Representative raw Western blot data is shown in Fig S6.

### OprF exists predominantly in the one-domain conformation in OMVs

Having established that OprF is equally distributed between OMVs and the OM, we were ready to test the hypothesis that OprF *conformation* differs between these two fractions. We accomplished this using a dot-blot analysis (Fig 6A). Purified OMVs and whole cells were deposited onto nitrocellulose membrane. The membrane was then probed using anti-V5 antibody to quantify the amount of V5 epitope exposed on the surface of the OMVs vs. cells. The amount of membrane (assessed as [KDO]) was titrated for each sample so that spots with equivalent antibody signal could be compared to avoid any issues with dynamic range in Western blots with widely varying amounts of signal (Fig 6B and Fig S7). The amount of membrane that needed to be added to achieve equal signal intensity was measured, allowing us to compare the relative amount of one-domain OprF present per unit membrane in OMVs *vs.* OM. Fig 6C shows that OMVs contain 10.5-fold more one-domain OprF than the cellular OM, while we previously established that the level of total OprF is equivalent in these two fractions. This result strongly suggests that the conformational distribution is flipped from nearly all two-domain OprF in the OM to nearly all one-domain OprF in OMVs.

**Fig 6.**
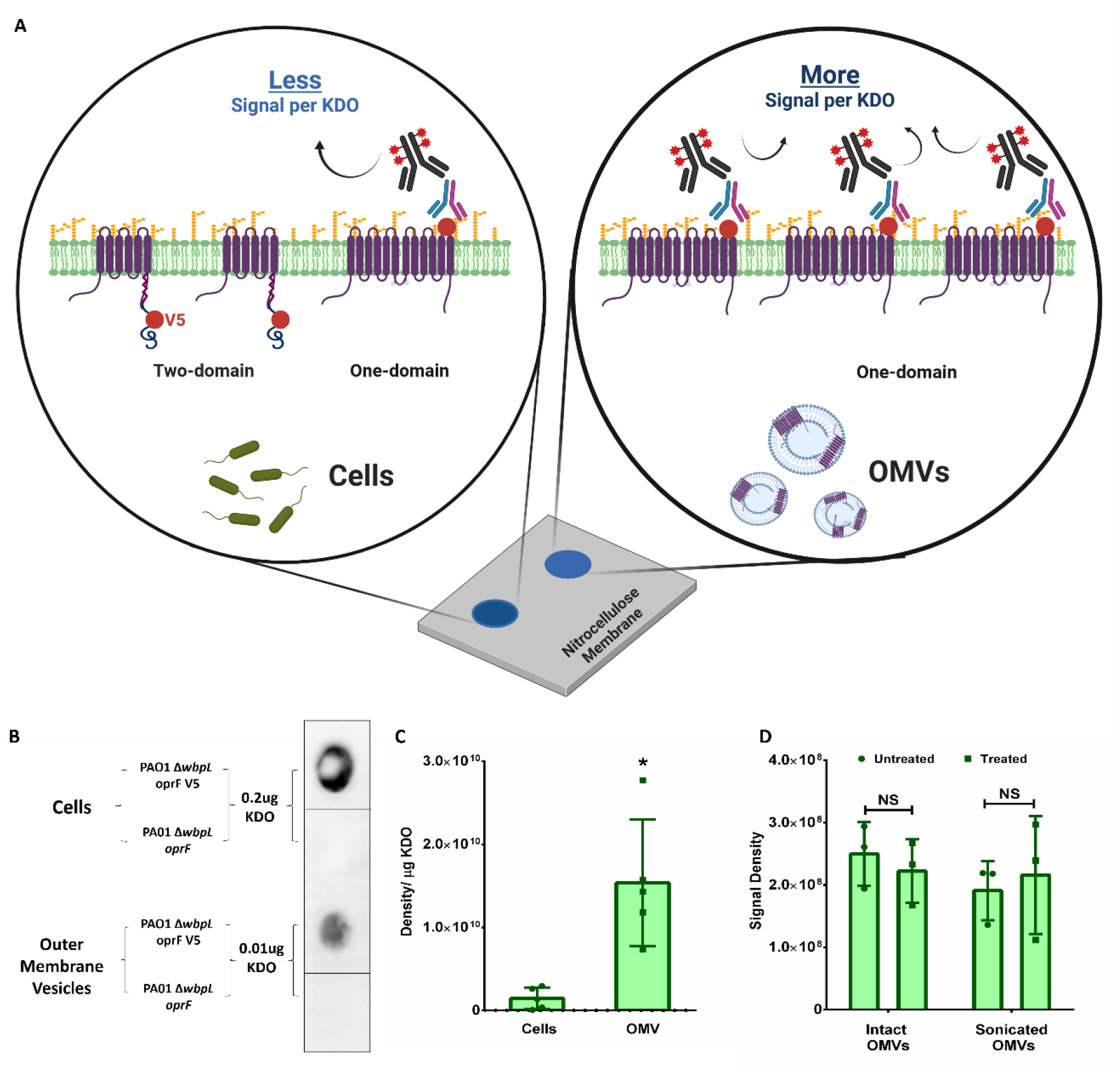
Analysis of OprF conformation in cells *vs*. OMVs. (A) Schematic representation of the dot-blot experiment used to assess OprF conformation in cells *vs*. OMVs. (B) Representative dot-blot showing samples of cells and OMVs deposited to reach comparable anti-V5 signal. The amount of total membrane spotted (0.2 ug *vs*. 0.01 ug KDO) is depicted (also see Fig S7). Negative controls expressing untagged OprF are also shown. (C) Normalized Density of anti-V5 signal per KDO for cells vs. OMVs from the dot-blot analysis. The level of accessibility of V5 epitope to exogenous antibody represents the amount of OprF present in the one-domain conformation. Error bars represent standard deviation. Statistical significance was analyzed by Two-tailed unpaired t-test with Welch’s correction (* p < 0.05; n = 5). (D) Proteolysis was performed on intact and sonicated OMVs to assess whether the C-terminus of OprF in OMVs is protected by insertion into the membrane (i.e. folded in the one-domain conformation). Loss of anti-V5 signal after Proteinase K treatment would represent degradation of the C-terminal domain. Error bars represent standard deviation. Statistical significance was analyzed by Multiple T-test which was corrected using Holm-Sidak method (NS = not significant; n = 3).

### Protease protection assay corroborates one-domain conformation of OprF in OMVs

Fig 6C demonstrates a dramatic difference in OprF conformation between OMVs and the OM. To further support our finding that the one-domain conformer predominates in OMVs, we performed a proteolysis protection assay on purified OMVs. The C-terminal domain of OprF is protected in the periplasm (and therefore the OMV lumen) when in the two-domain conformation but is susceptible to degradation if protease is allowed access (see Fig 1). Proteinase K treatment of intact OMVs would therefore be predicted to have no effect on the V5 epitope, which is present at position 312 in the C-terminal domain. In contrast, disruption of the OMVs by sonication while in the presence of proteinase K would give access to the lumen and facilitate degradation of the V5 epitope in any two-domain conformers present in the OMV sample. Fig 6D and Fig S8 show that protease treatment had no effect on the OprF-V5 signal in intact OMVs, as expected. Importantly, the V5 epitope was also fully protected in the sonicated OMV sample, supporting the proposal that the vast majority of OprF in OMVs is present in the one-domain conformation.

## Discussion

It is becoming clear that the ability to traffic cargo in extracellular vesicles is a feature common to all domains of life (reviewed here (88, 89)). In higher eukaryotes, there are several sophisticated protein-mediated pathways used to generate, package and traffic membrane vesicles. These systems incorporate familiar proteins such as clathrin, caveolin, dynamin, ESCRT proteins, SNAREs, etc. In archaea, it has also been proposed that ESCRT-related proteins play a role in extracellular vesicle formation (90, 91). Although dynamin-like proteins have recently been identified in some bacterial species (92, 93) and have been associated with secretion of proteins thought to be OMV cargo (94, 95), very little is known about whether proteins contribute specifically to OMV formation. A major question in the field, leading to several competing models (96, 97), has rather been to understand how bacteria could achieve vesicle budding in the absence of eukaryotic-like protein systems. We propose that prior to the evolution of sophisticated protein systems for membrane bending and scission, Gram-negative bacteria developed a program for vesicular transport based upon membrane curvature manipulation (governed by small molecule interactions (51) or lipid sorting (45, 47, 98)) in concert with a mechanism to selectively release patches of the OM from their underlying connections to PG. These “loosened” patches would then be free to bud under the influence of their own continued curvature induction or would be aided by periplasmic turgor (36, 99). With this work, we have uncovered a mechanism by which *P. aeruginosa* can selectively detach certain regions of the OM to be released as OMVs, providing the missing link that could unify many contributing inputs into a generalized model of OMV biogenesis across species.

The degree of connectivity between the OM and PG has long been implicated as a contributing factor to OMV formation (61–66, 68, 100, 101). Equally important, however, must be the *distribution* of OM-PG connections. The tethering of the OM in specific local regions must become loose enough for an OMV to form in that spot. It has previously been reported that OM-PG anchoring proteins are generally excluded from OMVs (69, 102), with the notable exception of members of the OmpA family (72, 77, 103–107). This suggests that OmpA family proteins are, at a minimum, present at the sites of OMV formation. Several reports also support the idea that OmpA-PG linkages are particularly important for OMV biogenesis in multiple species. Although deletion of nearly any major OM-PG anchoring protein results in hypervesiculation, Deatherage *et al.* observed in *Salmonella* that the hypervesiculation phenotype of an *ompA* mutant could not be complemented by overexpression of Braun’s lipoprotein (which provides the most abundant OM-PG linkages in *Salmonella*), hinting at a special role for OmpA in OMV formation (60). Wessel *et al.* (67) carried out a similar analysis in *P. aeruginosa* (where the OmpA homolog is OprF). Deletion of *oprF* resulted in the expected hypervesiculation, but the phenotype was rescued when PQS biosynthesis was simultaneously blocked (67). To us, this solidified the idea that hypervesiculation in *oprF*/*ompA* knockouts is not due to random membrane shedding, but rather is tied to the regulated biosynthesis of OMVs (which in *P. aeruginosa* is initiated by PQS through the bilayer-couple model (6, 51)). Further conclusions from that work are difficult to draw because of the confounding observation by Wessel *et al.* that their *oprF* mutant overproduced PQS (67), which we did not observe in our study.

Nevertheless, we aimed to exclude the possibility that the *oprF* hypervesiculation phenotype was due to general membrane instability caused by the loss of an abundant OM protein. Rather, we sought to demonstrate that it was the consequence of specific loss of OprF-PG connections. We expressed two OprF variants in the *oprF* mutant background to test their ability to rescue the hypervesiculation phenotype. The PG_del_ and C_del_ variants were designed to restore the presence of OprF in the OM but lack the ability to bind to PG, because of the removal of the specific PG-interacting motif (83, 108) and the entire C-terminal region, respectively. Both variants were confirmed to be expressed, correctly translocated to the OM, and correctly folded (in this and previous (81) work). We first showed that neither the *oprF* mutant nor any of the complemented strains showed increased susceptibility to deoxycholate challenge, confirming that membrane integrity was not compromised. This result is consistent with Deatherage *et al.* (60), who showed in *Salmonella* that *ompA* deletion did not affect membrane integrity, while *lpp* and *tol*-*pal* deletions had a negative effect. Following this, we demonstrated that neither OprF variant was able to restore OMV production back to WT levels. Vesicle production in the C_del_ variant strain was not different from that of the *oprF* mutant but, interestingly, the PG_del_ variant appeared to partially rescue OMV production. We attribute this to the fact that we deleted only 12 of the 18 amino acids identified as the PG-binding motif (83, 108) because topology studies (85) raised the possibility that full deletion might disrupt an integral membrane region involved in folding into the one-domain conformer. This decision may have resulted in the ability of the PG_del_ variant to retain weak PG binding affinity in the two-domain conformer. Because the native OprF expressed *in trans* was capable of full rescue, we conclude that the *oprF* hypervesiculation phenotype is due to the lost ability of the protein to interact with PG and not because of any indirect effect of deleting an abundant OM protein.

The importance of localized OprF-PG connections in OMV production is supported by our analysis of OMV size in the various strains. When OprF was prevented from attaching to PG there was an increase in both the number of OMVs produced and the mode of the OMV size distribution, suggesting that OMV size is controlled by the density of OM-PG connections. This is consistent with our previous observations when inducing OMV production in multiple species through exogenous addition of PQS (56). In that case, we unexpectedly found that PQS-induced OMVs in *E. coli*, *K. pneumoniae* and *P. mirabilis* were more similar in size to the producer species’ typical OMVs than those of *P. aeruginosa*. This led us to conclude that OMV size was controlled by a species-specific process, rather than the properties of the OMV inducer. The average density of OM-PG linkages likely varies from species to species. Variations in this spacing may explain both the species-specific OMV size distributions seen in the past (56) as well as the increased OMV size demonstrated here when OprF was prevented from anchoring to the PG. Deatherage *et al.* reported no OMV size increase in an *ompA* mutant in Salmonella (60). However, that work did not report average or mode OMV sizes and the size distribution histograms do appear to show a subtle bias toward larger OMVs in the relevant diameter ranges (<500nm) that would be consistent with what we report here.

If OM connectivity through OprF is indeed a control mechanism for OMV formation, as we propose, it would be predicted that OprF in naturally produced OMVs would predominantly exist in a PG-released form. A fascinating feature of OprF (and its homologs) is that they belong to the class of “slow porins” that have the ability to adopt both a two-domain (bound to PG in the periplasm) and one-domain (released from PG and inserted into the OM) conformation (79). We set out to answer the question of whether OprF serves as an OMV release latch by examining the conformational distribution of the protein in OMVs vs. the cellular OM.

To examine OprF conformation in purified OMVs and live cells, we expressed a variant OprF with a small epitope tag at a position that will only be exposed to the extracellular surface when the protein folds into the one-domain conformation (85). This allowed us to interrogate the amount of OprF in the one-domain conformation in each case using a simple dot blot. Before we could undertake this straightforward approach, it was necessary to understand whether OprF was sorted preferentially into OMVs and to establish a way to normalize the amount of OprF present in OMV vs. whole cell samples. By probing for the protein using Western blot and comparing that signal to the amount of LPS (KDO sugar) in each sample, we were able to confirm that the protein is equally sorted between OMVs and the OM.

With the understanding that the amount of OprF per unit membrane is equivalent in OMVs and cells, we went on to test the proportion of the protein present in the one-domain conformation in each case. The antibody signal was so starkly different between the two samples that we could not prepare the dot blots by applying an equal amount of membrane to each dot. Rather, we titrated the amount of sample in each dot to achieve equal antibody signal between OMVs and whole cells, then recorded how much membrane was required for each sample. The extremely low signal-to-KDO ratio of whole cells confirmed that the vast majority (>90%) (80, 82) of OprF in the OM of cells is in the PG-attached two-domain conformer. The signal-to-KDO ratio for the OMVs, on the other hand, was more than 10-fold increased vs. whole cells. This demonstrates a near total reversal of the conformational distribution of OprF in OMVs and establishes that the regions released as OMVs in native cells have very low OprF-PG connectivity. This idea is consistent with the recent observations of Orench-Rivera *et al.* (109). Though their analysis did not probe the involvement of the one- vs. two-domain conformations, these authors noted that when a truncated form of OmpA was expressed alongside full-length protein in *E. coli*, the truncated protein was more likely to be found in OMVs owing to its loss of PG contact. We confirmed the difference in OprF conformational distribution in native OMVs vs. whole cells using a protease protection assay. Disruption of the OM, allowing access of protease to the periplasm of cells, resulted in nearly complete degradation of the c-terminal domain in the >90% (80, 82) two-domain conformer found in cells. In contrast, providing protease access to the lumen of native OMVs had no effect on the c-terminal domain. This result solidifies the conclusion that OprF is found almost exclusively in the one-domain conformer in OMVs.

Our discovery that OprF is found in the “open” one-domain conformation in OMVs has several important consequences for OMV physiology and function that help to explain previous observations. The one-domain conformer folds as a non-specific porin, suggesting that OMVs would be much more permeable to small molecules than the cellular OM. This connects to a body of literature (110–112) demonstrating that OMVs are very effective at degrading antibiotics targeted at the producing bacteria (or if exogenously provided to other susceptible bacteria). This is true despite the fact that the antibiotic-degrading enzymes are protected *inside* the OMVs. Increased permeability owing to the dominant one-domain conformer in OMVs explains this. In the biofilm context, OMVs are known to be an abundant component of the matrix that encases cells (26, 29, 113–116). Following from the known interaction between cell-bound OprF and the lectin LecB (117), it has been speculated that an OprF-LecB-Psl tripartite interaction may stabilize OMVs in the matrix and retain them in the biofilm (118). Our finding that OprF exists in a very different conformation in OMVs vs. cells, however, introduces the possibility that OMV-resident OprF may interact with the matrix in a different manner than cells (or not at all). This would have major impacts on the mobility of OMVs in the matrix and their ability to transfer cargo between cells or even out of the biofilm altogether. In a similar vein, OMVs have long been believed to interact with eDNA in the biofilm matrix (119–121). Though no direct interaction with OprF has been proposed, loss of the protein was shown to affect overall eDNA production under some conditions (122). Thus, the role of OprF in OMV-matrix interaction is an area ripe for deeper investigation, and a new understanding that OprF in OMVs adopts a different conformation than OprF in cells provides a new wrinkle to be further explored.

Another intriguing question concerns *how* OprF comes to acquire such a different conformational distribution in OMVs vs. cells. Seminal work characterizing the two conformations of OprF identified four critical cysteine residues and proposed that each conformer arises from a dedicated folding pathway (80, 82). The interpretation of our results according to that model would be that OprF conformation is static, and the selective packaging of the one-domain conformer into OMVs would have to come from directed arrangement of these conformers in regions destined to become OMVs prior to initiation of OMV formation. In contrast, van der Heijden *et al.* observed that the OmpA transport pore in *Salmonella* could be rapidly opened or closed in response to oxidative stress (123) through a process that appeared to involve the reordering of disulfide linkages in the protein. This presents the exciting possibility that OprF (and homologs) could dynamically switch their conformation and thereby coordinate localized PG-release from regions of the membrane that otherwise would contain >90% two-domain conformers. This question continues to be an area of intense research in our laboratory.

With this work, we discovered a complete reversal of conformational distribution of OprF between the OM of cells and OMVs. This selective packaging is made possible by the unique ability of the OprF protein to adopt both a PG-bound and PG-unbound conformation. This discovery provides a mechanistic explanation for previous reports highlighting the particular importance of proteins like OprF in the vesicle formation process. Synthesizing this new information into what is already known about the process, we propose that OMV biogenesis proceeds through at least two distinct phases: (1) membrane curvature induction, possibly initiated in different ways under different conditions; and (2) OM release mediated by the removal of location-specific OprF-PG linkages. We dub this second phase the “Latch Model” of OMV biogenesis. Owing to the broad conservation of OmpA family (OprF homolog) proteins among bacteria, this model for later steps in OMV formation may serve as a common pathway to advance curvature events that were initiated by small molecule insertion (51), lipid remodeling (45, 47, 98), stress events (40, 99), etc. onward to the final stages of vesicle packaging and release. From a broader evolutionary perspective, this work also helps to shed light on how the ubiquitous process of extracellular vesicle formation can proceed using simple protein systems in the energy-poor environment of the bacterial periplasm.

## Materials and Methods

### Bacterial strains and general growth conditions

The bacterial strains and plasmids used are listed in Table 1. *P. aeruginosa* and *E. coli* cultures were routinely cultured in Lysogenic Miller broth/agar (Fisher Scientific). For all other purposes, the cultures were grown in Brain Heart Infusion (BHI) broth (Fisher Scientific). When necessary, media were supplemented with arabinose (Thermo) at 1% (w/v), gentamicin (Amresco) at 20 μg/mL for *E. coli* and 50 μg/mL for *P. aeruginosa* or with carbenicillin (IBI Scientific) at 250 μg/mL for *P. aeruginosa* strains.

### DNA Manipulations

Restriction endonucleases and T4 DNA ligase were purchased from New England Biolabs. DNeasy Blood & Tissue Kits (Qiagen) was used to isolate chromosomal DNA from *P. aeruginosa*. EconoSpin spin columns for DNA (Epoch Life Science) were used to isolate Plasmid DNA and genomic DNA fragments. DNA was amplified using Platinum *Taq* Polymerase High Fidelity (Invitrogen) or AccuPrime™ Pfx DNA Polymerase (Thermo Fischer Scientific).

### Construction of *oprF* PG_del_, *oprF* CT_del_ and *oprF* V5 mutants

The *oprF* PG_del_ construct was made by splicing overlap extension PCR (SOE PCR) (124) using PA14 genomic DNA and the SOE primers listed in Table 2. The *oprF* C_del_ construct was created using PA14 genomic DNA as template with the *oprF* forward and oprF C_del_ reverse primers (Table 2). To create *oprF* V5 construct, the codon for alanine (5’-GCT-3’) at position 312 in the protein sequence was replaced by the sequence encoding the V5 short tag in the reverse primer (Table 2). PAO1 genomic DNA was used as template. The final PCR products for *oprF* PG_del_, *oprF* CT_del_ and *oprF* V5 were purified and digested using XbaI and SacI restriction enzymes to allow for cloning into pMJT-1. Final sequences were verified using the Seq primers listed in Table 2. The plasmids were eventually purified from *E. coli* DH5α and electroporated into either the PA14 *oprF*::Tn, PAO1 Δ*oprF* or PAO1 Δ*wbpL* strains using methods described previously (125). Transformants were screened using LB agar containing 250 µg/mL carbenicillin.

### Purification of OMVs

Cultures of each strain were inoculated at starting OD_600_ of 0.01 into 30mL of fresh BHI medium supplemented with or without 250 µg/mL carbenicillin and 1% (w/v) arabinose for induction of the genes expressed through pMJT-1 plasmid. Cultures were incubated at 37°C with shaking (250rpm) until early stationary phase. Cells were separated from supernatant by centrifugation at 15,000×g for 15 minutes at 4°C (ThermoFisher Fiberlite F15s-8×50c rotor). The supernatant was collected, and any remaining cells were removed by filtration through a 0.45μm polyethersulfone (PES) syringe filter. The filtered supernatant was centrifuged at 209,438×g for 90 minutes at 4°C (Thermo S50-A rotor). The pelleted OMVs were resuspended in 1ml MV Buffer (50 mM Tris, 5 mM NaCl, 1 mM MgSO4, pH 7.4) (126).

### OMV concentration and size quantification

OMV number and size distribution were determined using the Malvern NanoSight NS300 instrument along with Nanoparticle Tracking and Analysis software (NTA 3.2) (31, 54, 56). To obtain 20 to 100 particles per frame (as per the manufacturer’s instructions), OMV samples were diluted 1:400 to 1:3200. The camera level was manually set to 12 with a gain of 1. Using different fields of view, each sample was analyzed three times for 30sec at 25°C. OMV concentration was normalized to OD_600_ of the extracted culture.

### Membrane stability assay

To assess whether the OprF variants caused membrane instability, we measured sensitivity to sodium deoxycholate (DOC), an ionic detergent, as previously described (60). Cultures were grown to late-exponential phase (OD = 0.5-1.0) and then plated on LB agar with carbenicillin (250µg/mL) ± 0.1% DOC (w/v). Plates were incubated overnight at 37°C and colony forming units were counted. Ability to withstand the presence of detergent was expressed in fold change when compared to growth without detergent.

### PQS quantification

PQS was quantified as previously described (126, 127). Briefly, PQS was extracted from bacterial cultures in 1:1 acidified ethyl acetate (0.1 mL/L acetic acid) before the organic phase (containing PQS) was removed and dried under nitrogen gas. Dried samples were resuspended in 25 µL optima grade methanol (Fisher) and spotted onto a straight-phase phosphate-impregnated TLC plate (EM Biosciences). PQS standards (Sigma) (100 µM-500 µM) were spotted on the same plate. The TLC was developed using 95:5 dichloromethane-methanol as mobile phase. PQS has intrinsic florescence which was excited under long-wave UV light and visualized (UVP Gel Doc-It2). Digital images were captured and analyzed using VisionWorks® LS Image Acquisition and Analysis software.

### Cell Fractionation

This protocol was adapted from our previously published work (54, 128). All the test strains were grown in 500mL BHI for 18hrs at 37°C and 250rpm. Cells were harvested by centrifugation at 15,000×*g* for 15 minutes at 4°C (ThermoFisher Fiberlite F15s-8×50c rotor). The cell pellets were washed in 100ml of 30mM Tris-HCl buffer (pH 8.0) and centrifuged again at 10,000×*g* for 10 minutes at 4°C. Washed pellets were then resuspended in 30mL of 20% Sucrose (prepared in 30mM Tris-HCl buffer, pH 8.0). The cells were lysed via sonication in the presence of 10 uL/mL HALT protease inhibitor cocktail plus EDTA (Thermo Fisher). Unlysed cells were removed by centrifugation at 10,000×*g* for 10 minutes. The supernatant was filtered through a 0.45µm syringe filter and the total membrane fraction was collected through ultracentrifugation at 100,000×*g* for 1 hour. The pellet obtained was resuspended in 1mL of 20% Sucrose (prepared in 30mM Tris-HCl buffer, pH 8.0). The supernatant was saved to be used as cytoplasmic fraction for downstream processes. To separate inner and outer membrane further, the pellet in 20% sucrose was layered on top of a sucrose gradient composed of 3mL of 70% (w/v) sucrose under 8mL of 60% (w/v) sucrose. The gradient was centrifuged at 32,700 rpm for 18h at 4°C (Beckman SW41-Ti rotor). After the centrifugation, 500µL fractions were collected from the top of the gradient. Sucrose concentration was reduced by diluting fractions into 30mM Tris-HCl buffer, pH 8.0 and samples were centrifuged at 100,000×*g* for 1hr (Thermo S120AT2 rotor). The washed pellets were resuspended and stored in 500µL of 30mM Tris-HCl buffer, pH 8.0.

### 3-deoxy-D-manno-2-octulosonic acid (KDO) assay

This protocol was modified from previous methods (129–131). 25µL of sample and KDO standards were mixed 1:1 with 0.5M H_2_SO_4_ and boiled for 15 min. After cooling at room temperature for 10 min, 25µL of 0.1 M periodic acid was added to each sample. Samples were vortexed and incubated at 25°C for 10 min after which 100µL of 0.2M sodium arsenate in 0.5M HCl was added. The samples were vortexed and 400µL of freshly prepared 0.6% thiobarbituric acid was added. Each sample was boiled for 10 min and then cooled at room temperature for 30-40 min. After the addition of 750µL of acidified n-butanol, the organic layer was recovered, and absorbance was measured at 552 nm and 509 nm. The measurements at 509 nm were subtracted from the measurements at 552 nm and the resulting values were compared to identically treated KDO standards.

### Succinate Dehydrogenase (SDH) assay

This protocol was adapted from Kasahara and Anraku (132). 200 uL of reaction mixture (50 mM Tris-HCl (pH 8.0), 4 mM potassium cyanide (KCN), 20 mM disodium succinate) was preincubated for 5 mins at room temperature in the wells of a 96-well plate. After this, 10 µL of sample was added to the mixture and further incubated for 5 mins at room temperature. To initiate the reaction, 2,6-dicholorphenolindophenol (DCPIP) and 0.2 mM phenazine methosulfate (PMS) was then added to the mixture and the SDH activity was measured by recording the absorbance at 600nm over time at 25°C (Tecan Infinite M-200 Pro).

### Immunoblotting analysis

0.5-1 µg total protein, based on Bradford assay (133), of the inner membrane (IM), Outer Membrane (OM) and Cytoplasmic (CP) fractions isolated from bacteria grown in BHI were separated by SDS-PAGE (12% gel). Proteins were transferred to a PVDF membrane (Bio-Rad Trans-Blot Turbo) and blocked in 5% skim milk in Tris Buffered Saline with 0.1% Tween-20. The membrane was then incubated with anti-OprF MA7-1 antibody (1:10000; kind gift of Dr. R.E.W. Hancock) (85). After washing, the membrane was incubated with HRP-conjugated donkey anti-mouse IgG antibody (1:1000) (Jackson Immuno Research), washed again, and exposed to Super Signal West Pico Chemiluminescent substrate (Thermo) (1:1000). Bands were visualized using the Gel Doc-It2 system (UVP) and quantified using VisionWorks® LS software.

### Protease Protection Assay

To determine the conformational state of the modified OprF PG_del_ protein, proteolysis of membrane-associated OprF was performed using a modification of the protocol of De Mot & Vanderleyden (83). Outer membrane fractions from PA14 *oprF*::Tn *oprF* PG_del_ and PAO1 Δ*oprF oprF* PG_del_ strains, prepared as described above, were suspended in 500 µL of 30 mM Tris-HCl, pH 8.0. Half of each sample was incubated with 100 ug/mL Proteinase K (Epoch Life Science) at pH 8.0 at 37°C for 30 mins. The reaction was stopped by adding PMSF at a final concentration of 0.3mg/mL. The other half of each sample was used as a control to compare with the protease-treated samples. Total protein was quantified in the treated and untreated samples using the Bradford assay. Proteolysis results were analyzed by Western blot as described above.

To determine the conformational state of OprF V5 in the vesicles, OMVs from PAO1 Δ*wbpL oprF* V5 were harvested as described above and split into four reactions of 100µL. The first two reaction mixtures served as “intact” OMV samples. One of these OMV samples was treated with 100 ug/mL proteinase K, while the other one functioned as an untreated control. The third and fourth reaction mixtures served as “sonicated” OMV samples. One of these OMV samples was incubated with proteinase K enzyme while being sonicated, in order to expose the lumen of the vesicles to the enzyme. The other sample was sonicated in the absence of Proteinase K. Proteinase K treatment was conducted at 37°C for 30 minutes, after which the reaction was stopped by adding trichloroacetic acid to a final concentration of 10% (w/v). Samples were diluted into 1 mL 30 mM Tris buffer (pH 8.0) and ultracentrifuged at 100,00*×*g for 1hr (S120-AT2 fixed angle rotor). The pellet was resuspended in 100 µL 30 mM Tris buffer (pH 8.0) and KDO assays were performed (as above) to analyze the total amount of outer membrane present in each sample. The samples were loaded based on equal amounts of KDO (0.2 µg/lane), separated by SDS-PAGE (12%), and analyzed by Western blot as described above (but with anti-V5 primary antibody).

### OprF sorting between OM and OMVs

PAO1 Δ*wbpL oprF* V5 was grown at 37°C (250 rpm) in 500 mL BHI, supplemented with 1% Arabinose and 250 µg/mL Carbenicillin, until early stationary phase. Cells were collected by centrifugation at 15,000xg for 15 minutes and the supernatant was used to harvest vesicles (described above) after being concentrated to ≤ 50mL by tangential flow filtration. Cells were lysed by sonication and membranes were fractionated as described above. The OM and OMV samples obtained were loaded based on equal amounts of KDO (0.2 µg/lane) and separated by SDS-PAGE (12%). The amount of OprF-V5 present in each fraction (normalized to KDO) was analyzed by Western blot using anti-V5 antibody as described above.

### Dot-blot assay

The conformational state of OprF-V5 in bacterial cells vs OMVs was determined by first collecting cells (described above) from 1 mL of PAO1 Δ*wbpL oprF* V5 culture, grown in 500 mL BHI supplemented with 1 % arabinose and 250 µg/mL carbenicillin until early stationary at 37°C (250 rpm). The cells were stored in the presence of HALT protease inhibitor cocktail plus EDTA (Thermo), at 4°C for downstream processing. The rest of the bacterial culture was used to harvest OMVs (described above). PAO1 Δ*wbpL oprF* bacterial cells and vesicles were harvested identically to be used as negative controls (no V5 epitope). KDO assay (described above) was used to determine the amount of outer membrane present in both cell and OMV samples. Bacterial cells and OMV samples (10 µl) with total 0.2 µg and 0.01 µg KDO, respectively, were then spotted on a pre-wetted (in Tris Buffered Saline with 0.1 % Tween-20) nitrocellulose membrane and dried for 1 hour at RT. The membrane was then blocked with 5% skim milk in Tris Buffered Saline with 0.1% Tween-20. The Dot-blot was visualized using anti-V5 antibody (Thermo), and HRP-conjugated donkey anti-mouse IgG antibody (Jackson Immuno Research) followed by chemiluminescence analysis as described above. The antibody signal obtained from PAO1 Δ*wbpL oprF* bacterial cells and vesicles (negative controls) was subtracted from the signal obtained from PAO1 Δ*wbpL oprF* V5. The final signal density from bacterial cells and OMV samples was then normalized to the amount of KDO present in each sample.

## Supporting information

Supplement (all)

## Acknowledgments

We thank Dr. Marvin Whiteley, Dr. Catalina Florez and Dr. Adam Cooke for their critical reading and insightful discussions about the manuscript.

This work was supported in part by a grant from the NIH (1R15GM135862 to J.W.S). Figure 6A was created using Biorender.com.

The content is solely the responsibility of the authors and does not necessarily represent the official views of the National Institutes of Health.

