## Supplement (all) for "OprF functions as a latch to direct Outer Membrane Vesicle release in *Pseudomonas aeruginosa*"

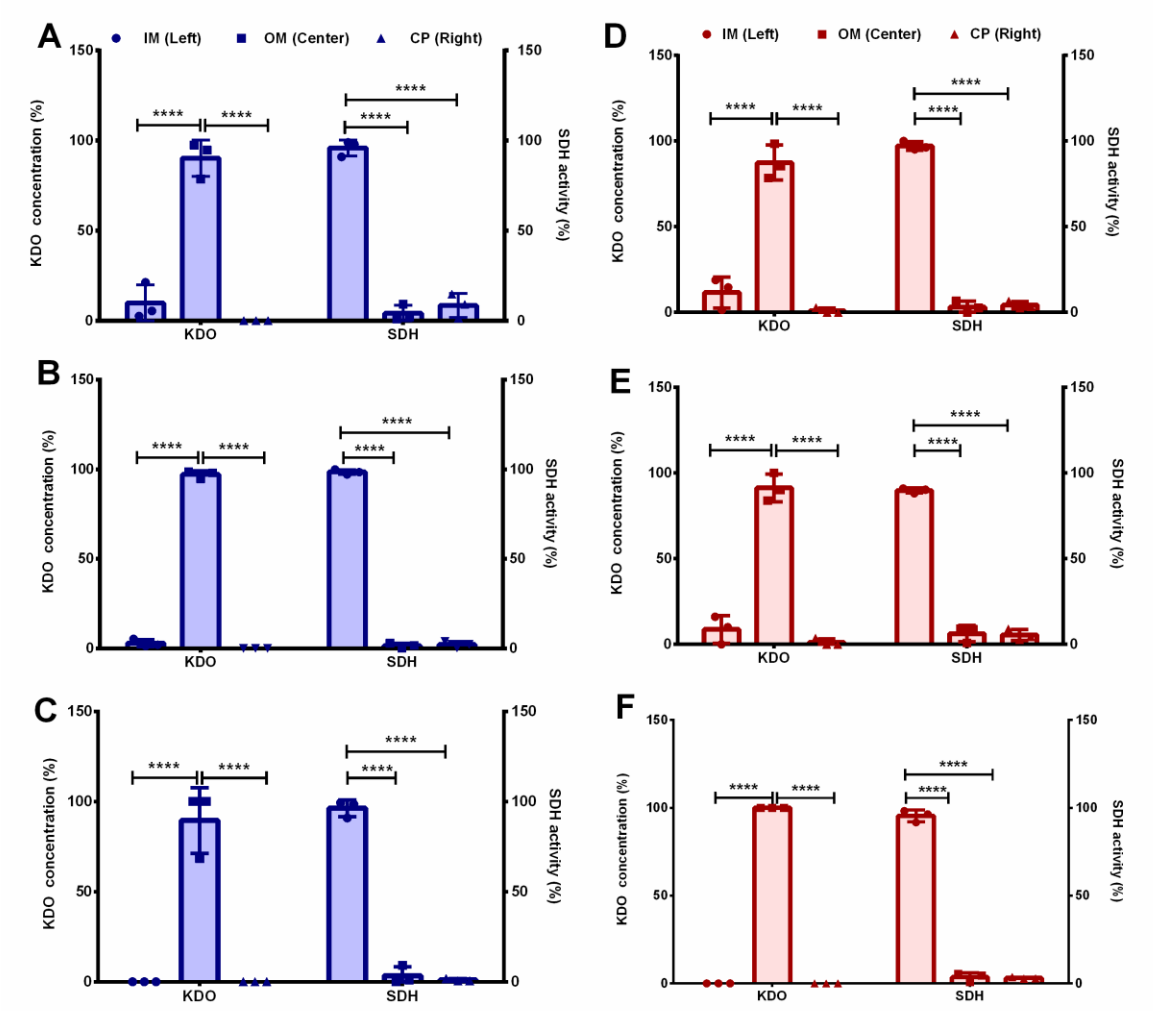


**Fig. S1.** Confirmation of separation of sub-cellular fractions**.** KDO and SDH assay on inner membrane (IM), outer membrane (OM), and cytoplasmic fractions (CP) of (A) PA14 *oprF* complement, (B) PA14 *oprF* $PG_{del}$ variant, (C) PA14 *oprF* $C_{del}$ variant (D) PAO1 *oprF* complement, (E) PAO1 *oprF* $PG_{del}$ variant, and (F) PAO1 *oprF* $C_{del}$ variant. Error bars represent standard deviation. Statistical significance was analyzed by Two-way ANOVA followed by Sidak's multiple comparisons test between IM, OM, and CP (****p < 0.0001) and n = 3.

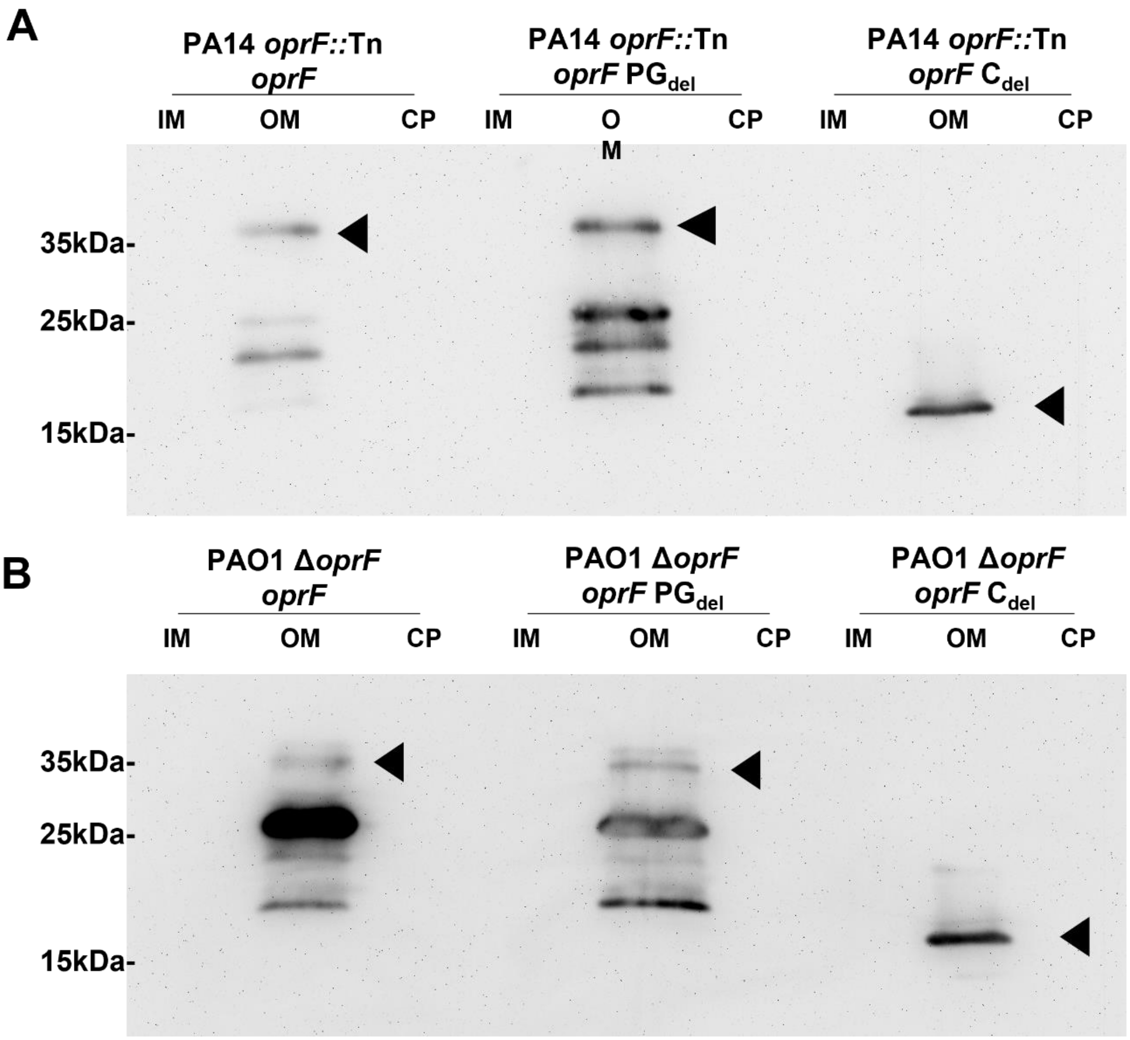


**Fig. S2.** Western Blot of OprF in different fractions. Western blot showing OprF (protein marked by black arrows) detected by antibody specific for the N-terminus region of the protein in outer membrane (OM), inner membrane (IM) and cytoplasmic (CP) fractions collected from (A) PA14 *oprF* complement, PA14 *oprF* $PG_{del}$ and PA14 *oprF* $C_{del}$ variants and (B) PAO1 *oprF* complement, PAO1 *oprF* $PG_{del}$, and PAO1 *oprF* $C_{del}$ variants. The figures are representative of three independent experiments.

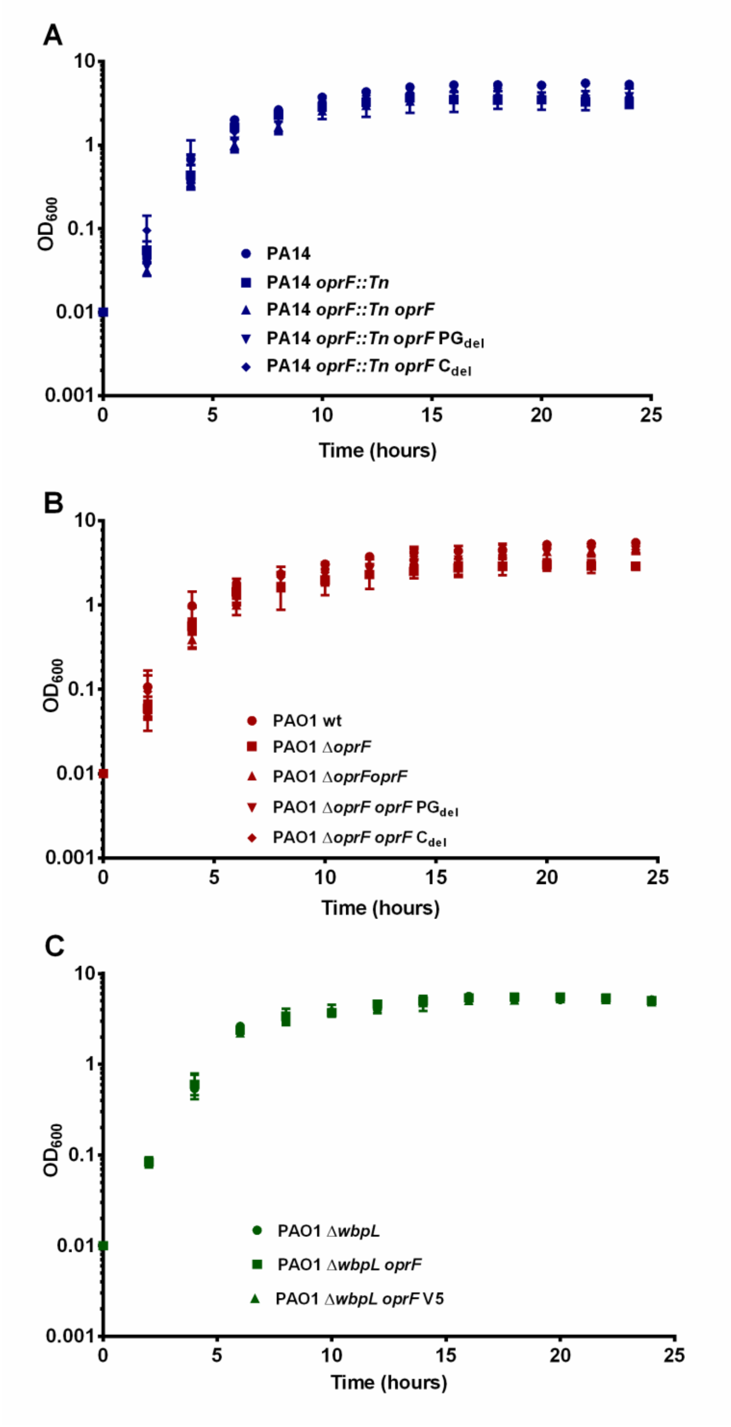


**Fig. S3.** Growth curve. The optical density measured over time for all the strains in (A) PA14 and (B) PAO1 and (C) PAO1 Δ*wbpL* background were plotted as mean ± standard deviation (n = 3), which were grown in BHI with shaking (250rpm) at 37°C.

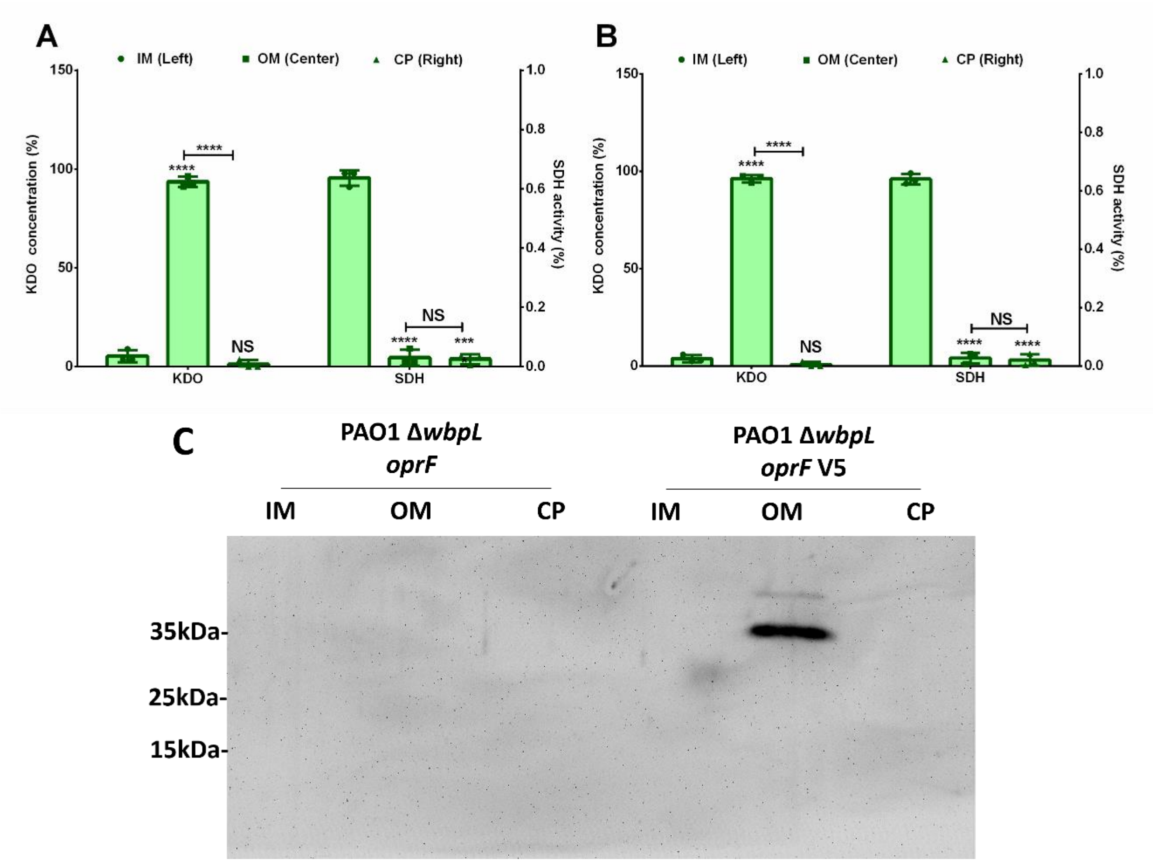


**Fig. S4.** Confirmation of membrane separation and presence of OprF-V5 in outer membrane. Separation of outer membrane (OM), inner membrane (IM) and cytoplasmic fractions (CP) was confirmed for (A) PAO1 Δ*wbpL* oprF and (B) PAO1 Δ*wbpL oprF V5* by measuring the presence of KDO sugar and SDH enzyme activity. Error bars represent standard deviation. Statistical significance was analyzed by Two-way ANNOVA followed by Sidak's multiple comparisons test between IM, OM, and CP (****$p < 0.0001$, NS= not significant) and n = 3. (C) Western blot using anti-V5 antibody on inner membrane (IM), outer membrane (OM) and cytoplasmic (CP) fraction from PAO1 Δ*wbpL oprF* and PAO1 Δ*wbpL oprF* V5. The figure is a representative of three independent experiments. Asterisks directly over bars denote differences from IM; asterisks associated with horizontal lines refer to differences between the indicated bars.

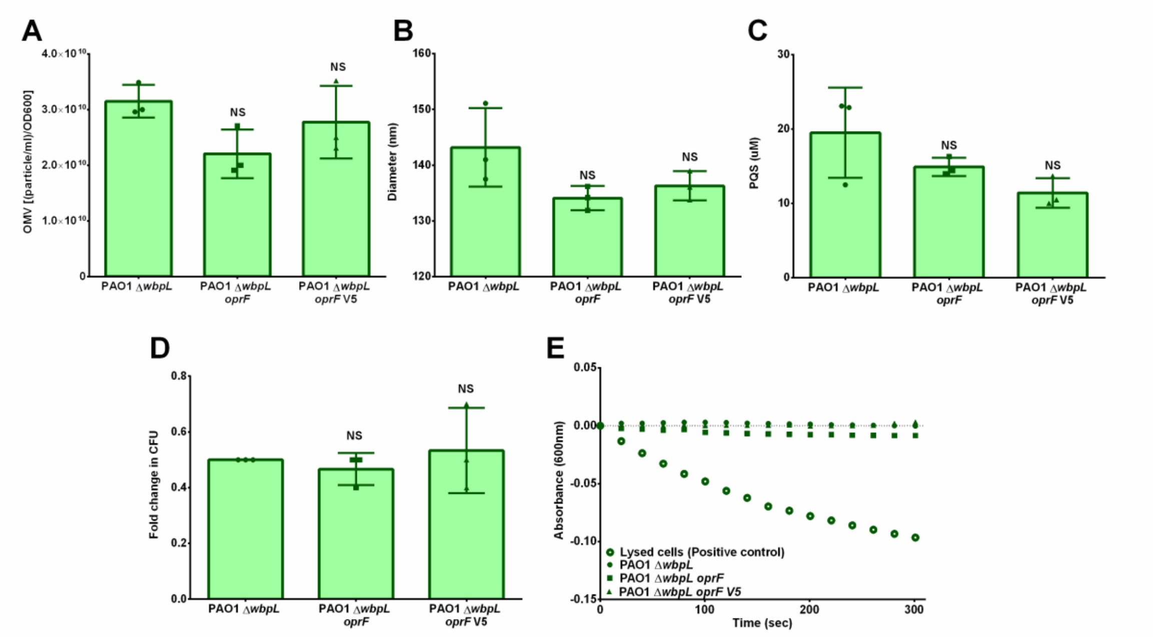


**Fig. S5.** Quality control on all the PAO1 ΔwbpL strains. Comparison of OMV (A) quantity and (B) size produced by all the strains tested. The OMV particle number and mode diameter was quantified using Nanoparticle Tracking and Analysis software (NTA 3.2). All strains were grown to early stationary phase. Error bars represent standard deviation. Statistical significance was analyzed by One-way ANOVA followed by Dunnett's multiple comparisons test and n = 3 (NS = not significant). (C) PQS was extracted and quantified from each sample using thin layer chromatography and densitometry analysis. Error bars represent standard deviation. Statistical significance was analyzed by One-way ANOVA followed by Dunnett's multiple comparisons test and n = 3 (NS = not significant). (D) Membrane stability was tested for all the strains when grown in presence of 0.1 % sodium deoxycholate. Error bars represent standard deviation. Statistical significance was analyzed by One-way ANOVA followed by Dunnett's multiple comparisons test and n = 3 (NS= not significant). (E) Succinate dehydrogenase enzyme activity was measured over time in the OMVs from all the strains and compared to lysed bacterial cells (positive control. The figure is a representative of three independent experiments.

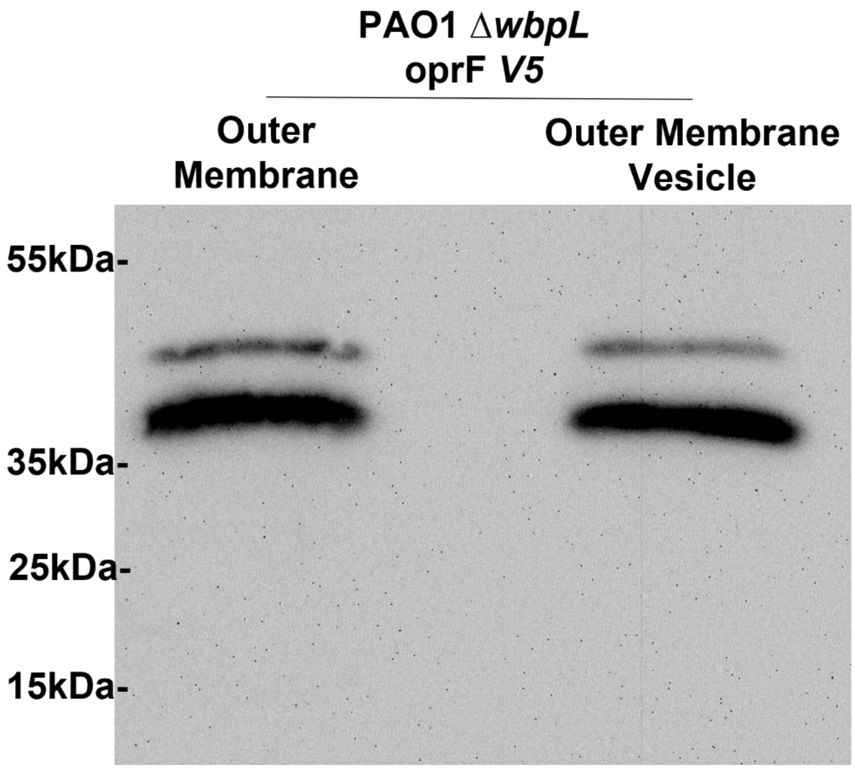


**Fig. S6.** Outer membrane (OM) fractions and vesicles harvested from PAO1 Δ*wbpL* V5 were loaded and separated on a 12 % SDS-PAGE gel based on equal amount of KDO (0.2 µg/lane) and transferred onto PVDF membrane. Western blots were carried out using anti-V5 antibody and two species of V5 tagged OprF, native (38.5 kDa) and mature (36.1 kDa) were detected in both OM and OMV samples from PAO1 Δ*wbpL* oprF V5. The figure is a representative of five independent experiments.

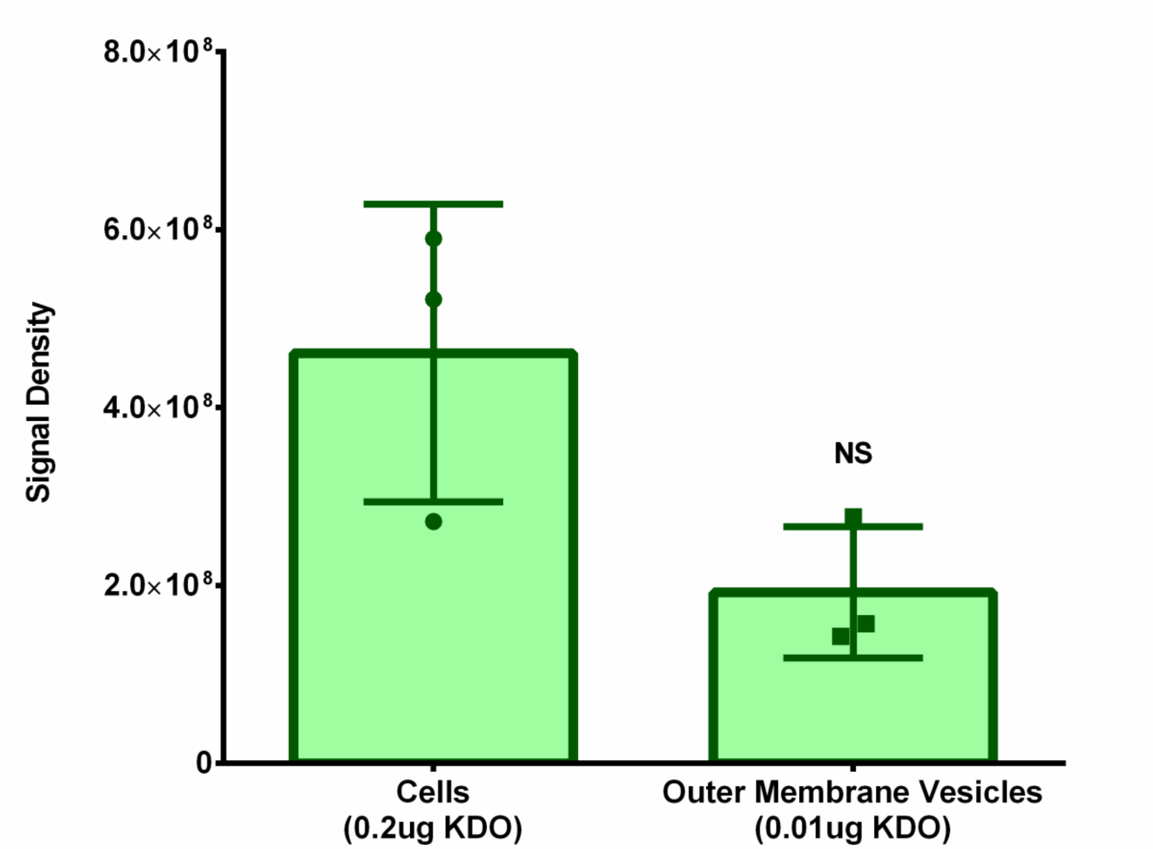


**Fig. S7.** Densitometry analysis after incubation with anti-V5 antibody of dot-blots shown in Fig 6B showing comparable signal density between cells and OMVs spotted based on total KDO of 0.2 µg and 0.01 µg per spot, respectively. Error bars represent standard deviation. Statistical significance was analyzed by Two- tailed unpaired t-test with Welch's correction. (NS=not significant) and n=3.

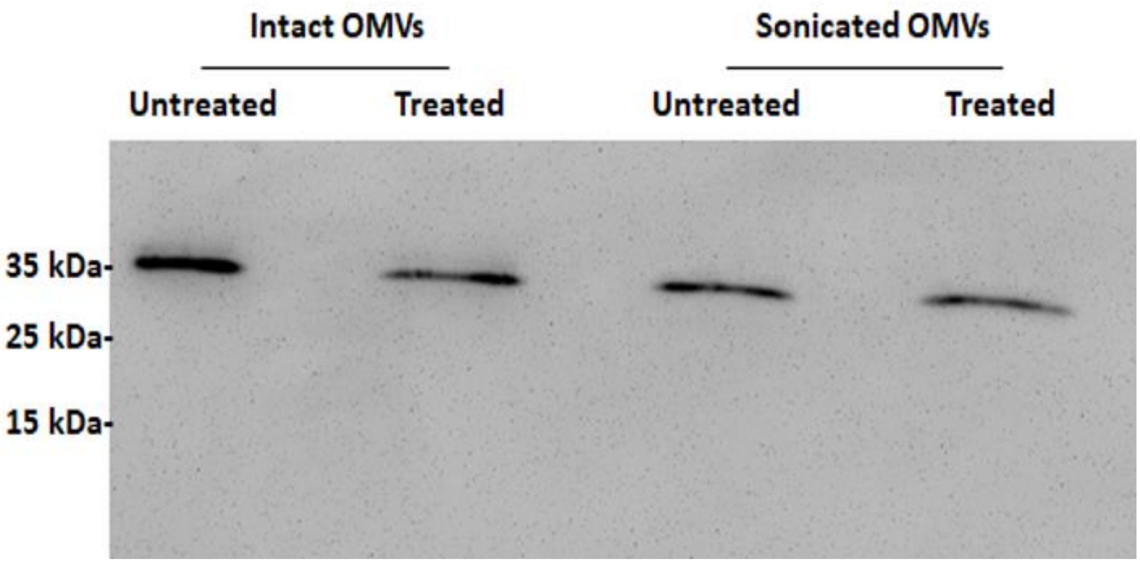


**Fig. S8.** Western blot using anti-V5 antibody on intact and sonicated OMVs which were either not treated or treated with proteinase K enzyme. All samples were loaded based on equal amount of outer membrane (KDO per lane = 0.02 µg). The image is representative of three independent experiments.

**Table 1. Strains and Plasmids used in this study.**

| Strain or Plasmid | Description | Reference |
|---|---|---|
| **Strain** | | |
| ***Escherichia coli*** | | |
| DH5α | F– Φ80lacZΔM15 Δ(lacZYA-argF) U169 recA1 endA1 hsdR17 (rK–, mK+) phoA supE44 λ– thi-1 gyrA96 relA1 | (86) |
| ***Pseudomonas aeruginosa*** | | |
| PA14 | Wild type *Pseudomonas aeruginosa* strain | (48) |
| *oprF*::*Tn* | PA14 *oprF::Mar2xT7* ($Gm^r$) | (48); kind gift from Dr. Karin Sauer |
| *P. aeruginosa* MPA01 | Wild type *Pseudomonas aeruginosa* strain | (87, 88); kind gift from Dr. Karin Sauer |
| ∆*oprF* | *oprF* clean deletion in MPAO1 background | kind gift from Dr. Boo Shan Tseng |
| PA01 ∆*wbpL* | Wild type *Pseudomonas aeruginosa* strain | (50, 51); kind gift from Dr. J.S. Lam |
| **Plasmids** | | |

| Strain or Plasmid | Description | Reference |
|---|---|---|
| pMJT-1 | $Carb^R$; *araC-pBAD* expression vector | (89) |
| pMJT-1 *oprF* | $Carb^R$; pMJT-1 derived *oprF* over expression vector | This study |
| pMJT-1 *oprF* $PG_{del}$ | $Carb^R$; pMJT-1 derived *oprF* $N_{265}$-$V_{276}$ *del* over expression vector | This study |
| pMJT-1 *oprF* $CT_{del}$ | $Carb^R$; pMJT-1 derived *oprF* $K_{164}$-$K_{326}$ *del* over expression vector | This study |
| pMJT-1 *oprF* V5 | $Carb^R$; pMJT-1 derived *oprF* $A_{312}$*IPNPLLGLD* over expression vector | This study |

**Table 2. List of Primers used in this study.**

| Oligonucleotide name | Sequence (5'–3') [a] |
|---|---|
| oprF Forward | CGCGCGC*TCTAGA***GACCATCAAGATGGGGATTTAACGGATG** |
| oprF Reverse | GGGGGGGG*GAGCTC***CGATTACTTGGCTTCAGCTTC** |
| oprF PG$_{del}$ SOE 5' | TCGTTGACCAGTACGTCACG**GTAAGCGTCGGTGCCGACG** |
| oprF PG$_{del}$ SOE 3' | CGTCGGCACCGACGCTTAC**CGTGACGTACTGGTCAACGA** |
| *oprF* CT$_{del}$ Reverse | GTCAGTCA*GAGCTC*TTA**CGAACCACCGAAGTTGAAGCCGA** |
| *oprF* V5 Reverse | GCAGTCA*GAGCTC***TTACTTGGCTTCAGCTTCTACTTCGGCTTCAACGCGACGGTTGAT**ATCCAGGCCCAGCAGCGGGTTCGGGAT**GCGGCCTTCAGCGG** |
| pMJT-1 Seq Forward | **GACCGCGAATGGTGAG** |
| pMJT-1 Seq Reverse | **GAGCTGATACCGCTCG** |

[a] Italics represent regions of enzyme restriction sites. Nucleotides in bold are regions on oligos which allow them to act as PCR primers on appropriate template.
